# Global effects of the guanine quadruplex binding ligands Pyridostatin and TMPyP4 on microRNA regulation in response to ionizing radiation in HeLa cells

**DOI:** 10.64898/2026.11.11.766052

**Authors:** Shruti Mishra, Himani Tewari, Sindhuja Singh, Swathi Kota

## Abstract

Guanine quadruplexes (G4s) are secondary structures in DNA and RNA that regulate a wide range of biological processes. Small molecules that assist in formation, stabilization or destabilization of G4s indirectly alter the cellular functions governed by these structures. Pyridostatin (PDS) and TMPyP4 are two well-characterized G4-binding ligands that have been shown to modulate various cellular functions, particularly in cancer cells. However, their effects on microRNAs (miRNAs), which are key regulators of post-transcriptional gene expression, remain poorly understood. HeLa cells treated either with PDS or TMPyP4 and gamma radiation were subjected to small RNA sequencing. Analysis results revealed a higher overall small RNA content in PDS treated cells, whereas TMPyP4 treated cells exhibited a greater number of differentially expressed miRNAs following gamma radiation exposure. Target gene analysis of the top differentially regulated miRNAs revealed that molecular functions related to DNA/nucleic acid binding activities of various factors, including RNA polymerase II cis-regulatory region sequence-specific binding activity, were largely affected. Furthermore, KEGG pathway enrichment studies suggested that observed changes may result from alterations in signal transduction pathways in cells treated with G4 ligands and gamma radiation compared to untreated control cells. Additionally, G4 ligand treatment altered the cellular expression levels of histone modifying enzymes, including histone deacetylases and methyltransferases. This finding suggest that the two G4 binding ligands-PDS and TMPyP4 differentially regulate miRNA expression profiles, histone modifying enzymes and associated biological pathways in HeLa cells.

## 1. Introduction

Guanine quadruplexes (G4s) are non-canonical secondary structures formed in nucleic acids containing continuous stretches of guanines. Four guanine bases connected by Hoogsteen hydrogen bonding form a square planar structure called G-tetrad. Stacking of these G-tetrad on top of each other results in the formation of complex four stranded guanine quadruplex structures. Monovalent cations such as potassium (K^+^) or sodium (Na^+^) coordinate with the electronegative oxygen atoms of the guanine bases to stabilize the G-tetrads (Spiegel J et al., 2020). G4s can be formed in both DNA and RNA. Depending on the strand orientation and folding patterns, DNA G4 structures can adopt parallel, hybrid or antiparallel topologies. In contrast, RNA G4s primarily adopt parallel topology because of the steric hindrance of 2’-hydroxyl group. RNA G4s are generally more stable than their DNA counterparts due to more intramolecular interactions and structural compaction (Fay MM et al., 2017). Since G4s discovery, DNA G4s have been extensively studied, leading to the establishment of their regulatory role in a variety of cellular processes such as DNA replication, transcription and maintenance of genome stability in both prokaryotes and eukaryotes (Mishra et al., 2021). Additionally, bioinformatics predictions and experimental evidences have shown that intracellular RNA G4s are involved in multiple biological functions including regulation of RNA stability, RNA maturation and translation processes. RNA G4s have been identified within the protein-coding transcripts, 5’ and 3’ untranslated regions (UTRs), non-coding RNAs and telomeric repeat-containing RNAs (TERRAs) (Kharel P and Ivanov P., 2024).

MicroRNAs (miRNAs) are a class of endogenously generated, non-coding RNAs that play a crucial role in the regulation of gene expression. Most miRNAs are transcribed by RNA polymerase II, while a smaller subset is transcribed by RNA polymerase III. The initial RNA transcript generated from DNA is known as the primary miRNA (pri-miRNA), which is processed in the nucleus by the Drosha complex to produce smaller intermediate molecules called precursor miRNAs (pre-miRNAs). These pre-miRNAs are further processed in the cytoplasm by the enzyme Dicer into short, mature miRNAs that are typically 21-23 nucleotide in length. Usually, mature miRNAs bind at the 3’ region of the target messenger RNA (mRNA), leading to translational repression or its degradation. A single mature miRNA may have multiple target mRNAs, and conversely, an individual mRNA can be targeted by many miRNAs, thereby forming complex regulatory networks (Finnegan E. F. and Pasquinelli A. E., 2013) As miRNAs are crucial regulators of cellular gene expression, their biogenesis and functioning are tightly controlled at multiple levels. Transcription of pri-miRNA is majorly governed by transcription factors and RNA polymerase II machinery, similar to protein-coding genes. The processing of pri-miRNA depends on its secondary structures as well as various co-factors like DGCR8, a component of the Drosha complex. Similarly, the Dicer mediated processing is modulated by RNA-binding proteins, signalling pathways and structure of pre-miRNAs itself (Ha M and Kim VN., 2014). Dysregulation of miRNA biogenesis or expression has been linked to the development of several pathological conditions such as neurodegenerative diseases, autoimmune/inflammatory diseases, cardiovascular diseases and cancer (Ha TY, 2011; Kapplingattu SV et al., 2025).

The role of G4s in the regulation of miRNA biogenesis and function has only recently begun to be explored. For instance, it has been shown that the formation of G4s within pri-miRNAs, near Drosha cutting sites, may affect Drosha binding and processing efficiency leading to change in miRNAs expression levels (Rouleau SG et al., 2018). In certain cases, stable G4s may also compete with the canonical stem-loop structure necessary for miRNA maturation as demonstrated in zebrafish embryos (Steeman TJ et al., 2023). In addition, an essential role of a conserved intramolecular G4 in biogenesis of specific miRNAs involved in cardiac function has been reported (Zhu M et al., 2021). Beyond miRNA biogenesis, G4s present within the 3’ UTR of target mRNAs can also modulate mRNA functions by regulating miRNA binding and target accessibility (Rouleau S et al., 2017). Although all these studies collectively show the regulatory part of G4s in miRNA expression, there are nil or very few reports on how intracellular G4s dynamics could influence the global expression profile of cellular miRNAs.

Here, we report changes in miRNA expression profiles in HeLa cells following treatment with two most studied G4 binding ligands, Pyridostatin (PDS) and 5,10,15,20-tetrakis-(*N*-methyl-4-pyridyl) porphyrin (TMPyP4) (Anantha NV et al., 1998; Hänsel-Hertsch R et al., 2017). HeLa cells treated with PDS and TMPyP4 showed differential cellular responses to gamma radiation induced DNA damage compared with untreated control cells. Several known and novel miRNAs were found to be differentially expressed in response to gamma radiation and G4 binding ligands treatment. Bioinformatics analysis revealed that the altered miRNA expression majorly affected molecular functions related to DNA binding proteins like transcription factors as well as cellular pathways involving signal transduction. In addition, some of the identified miRNAs were found to contain putative G4 forming motifs within their pre-miRNAs and fold into stable parallel G4 structures in vitro. PDS and TMPyP4 treatment also altered the expression of histone modifying enzymes, particularly deacetylases and methyl transferases. Altogether these results suggest that cellular G4s levels play an important role in the regulation of miRNA expression and chromatin dynamics.

## 2. Methodology

### 2.1 Materials

Dulbecco’s Modified Eagle Medium (DMEM), antibiotic–antimycotic solution, fetal bovine serum (FBS), dimethyl sulfoxide (DMSO) and trypsin were purchased from Himedia Laboratories Pvt. Lt. Anti-DNA G-quadruplex (G4) antibody clone 1H6 (MABE1126) were obtained from Sigma-Aldrich, St. Louis, USA and γ-H2AX antibody was procured from Cell Signaling Technology (9718S). Pyridostatin (PDS), 5,10,15,20-tetrakis-(*N*-methyl-4-pyridyl) porphyrin (TMPyP4), Thioflavin T (ThT) dye 2-(4-(Dimethylamino) phenyl)-3,6-dimethylbenzo[d]thiazol-3-iumchloride were procured from Sigma-Aldrich.

### 2.2 Cell culture

Human cervical cancer cell line - HeLa (CCL-2) was procured from National Centre for Cell Science (NCCS), Pune, India and maintained in cell culture media-DMEM supplemented with 10% heat-inactivated FBS, and 1% antibiotic–antimycotic solution at 37°C in a humidified atmosphere of 5% CO_2_. Cells were grown in 90-cm dishes and routinely passaged every 3–4 days when 70–80% confluent.

### 2.3 Cell treatment and irradiation

The cells were irradiated using Bhabhatron-II Telecobalt unit (Bhabha Atomic Research Centre, Mumbai, India) with ^60^Co having dose rate of 1 Gy/min at ambient temperature. The samples were irradiated at different single doses of 4 or 8 Gy. The samples were irradiated in 6-well plates or 90-cm culture dishes and cells were recovered after irradiation at the indicated time point of 30 min and 2 hr. In G4 binding ligand treated cells, PDS (5µM) and TMPyP4 (10µM) were added to the culture medium at 24 h before exposure to irradiation and maintained in the medium till the recovery period before cell collection or fixation.

### 2.4 Immunofluorescence and cell imaging

Cells were grown to 70-80% confluency on coverslips which were kept in 6-well plates and irradiated as mentioned above. Cell layer was washed at different time periods after irradiation in PBS and fixed for 20 min in 4% paraformaldehyde at room temperature. Afterwards, the cells were washed twice in PBS. Cells were permeabilized in 0.25%TritonX-100 in PBS for 5 min, washed two times in PBS and incubated with primary antibodies (G4 and γ-H2AX, 1:250 ratio) in blocking solution consisting 2% BSA in PBS. Cells were incubated with primary antibodies overnight at 4°C, after incubation the cells were washed three times in PBS and incubated with secondary antibodies anti-mouse IgG conjugated with Alexa Fluor 594 (Sigma) and Alexa Fluor 488 (Sigma) for 2 h in a blocking solution. Cells were again washed and the coverslips were mounted on slide with antifade DAPI mounting media. Cells were visualized under confocal laser scanning microscope (Olympus IX83 inverted confocal microscope) and the number of foci in ~300 cells were analysed using automated Olympus CellSens software and plotted using GraphPad Prism 8 software.

### 2.5 RNA isolation, library preparation and sequencing

RNA was extracted from HeLa cells and G4 ligand (PDS and TMPyP4) treated cells using Direct-zol RNA mini kit (Zymo Research). The frozen cells were resuspended with 1 mL of Trizol, mixed with equal volume of ethanol (100%) and loaded into Zymo-Spin IIICG Column placed in collection tube. The tubes were centrifuged at >10,000 rpm for 1 min, and flow through discarded. On column DNase I treatment and subsequent column washes were performed according to manufacturer’s instruction. The RNA was eluted from the column and the concentration and purity of RNA was quantified using the Nanodrop Spectrophotometer (Thermo Scientific; 2000). The percentage of miRNA in the samples was obtained on Agilent Bioanalyzer. The RNA concentration was quantified using Qubit RNA HS assay kit (Q32855). Small RNA sequencing (smRNA) libraries were constructed using QIAseq® miRNA Library Kit protocol (Cat: 331502; Qiagen, Maryland, U.S.A.). Prepared Illumina-compatible libraries were quantified using a Qubit fluorometer and verified for size distribution on an Agilent 2200 TapeStation. The libraries were subsequently pooled and sequenced on an Illumina NovaSeq 6000 platform using single-end 50 bp (SE50) sequencing chemistry. The data obtained from the sequencing run was demultiplexed using Bcl2fastq softwarev2.20 and FastQ files were generated based on the unique dual barcode sequences. The sequencing quality was assessed using FastQC v0.11.8 software.

The adapter sequences were trimmed and bases above Q30 were considered. The raw reads were then filtered to obtain specific length of 16-40 bases, which were subsequently aligned to genome using Bowtie to exclude RNA species such as r, t, sn and snoRNA. The unaligned reads were further analysed to predict known miRNAs using BLAST and novel miRNAs using Mireap. Differential gene expression analysis among various comparison groups was performed using DESeq. The target miRNAs were predicted using TargetScanHuman analysis (transcripts) for top 5 upregulated and downregulated miRNAs. miRNA-mRNA interaction networks of different comparison groups were generated using Cytoscape tool. Volcano plots were generated for the differentially expressed gene list using their normalized expression levels. Gene Ontology (GO) enrichment analysis was carried out with the human background database and a false discovery rate (FDR) cutoff of 0.05 using ShinyGO gene-set enrichment tool (http://bioinformatics.sdstate.edu/go, version 0.80). Secondary structures of pre-miRNA were generated using RNAfold webserver (http://rna.tbi.univie.ac.at/cgi-bin/RNAWebSuite/RNAfold.cgi).

### 2.6 In vitro validation of putative G4 motifs

The 5′ and 3′ untranslated regions (UTRs) of histone deacetylases (HDAC1 & HDAC2) and methyl transferases (SETD7 & SUV39H1) were retrieved using www.alliancegenome.org and analysed for the prediction of putative G4 forming sequence (PQS) using QGRS (http://bioinformatics.ramapo.edu/QGRS/index.php). The formation of stable G4 structures by few of the sequences of histone deacetylases (HDAC1 & HDAC2) and methyl transferases (SETD7 & SUV39H1) were checked by performing various in vitro assays The synthesized PQS (8 μM) were heated at 95°C in the presence of 50 mM KCl and 10 mM Tris buffer (pH 7.4) and slowly cooled down to room temperature. These annealed sequences were further used to perform the in vitro validation. The annealed sequences were mixed with final concentration of 0.5 μM Thioflavin T (ThT) in a 96-well microplate. The fluorescence spectra were collected between 440 and 700nm after excitation at 425 nm for ThT in a Synergy H1 Hybrid multi-mode microplate reader. The fluorescence intensity of the ThT alone or in the presence of putative G4 motifs were compared. The fluorescence emission spectra were plotted using GraphPad Prism 8.0.

Circular dichroism (CD) spectroscopy was also performed to examine the putative G4 DNA topologies in vitro, as discussed earlier (Tewari et al., 2025). CD spectra of annealed sequences were recorded on a spectrophotometer (Biologic MOS-500 CD spectropolarimeter), in the wavelength range from 220 to 320 nm with a scanning speed of 100 nm/min and a response time of 2s, using a quartz cuvette with a path length of 1.0 mm. All measurements were performed in triplicate, with the buffer baseline subtracted by the instrument’s software.

## 3. Results

### 3.1 Pyridostatin and TMPyP4 treatments show differential effect on the small RNA expression in HeLa cells

Small-molecule ligands such as Pyridostatin (PDS) and 5,10,15,20-tetrakis-(N-methyl-4-pyridyl) porphyrin (TMPyP4) are known to increase the frequency of intracellular G4s by binding to and stabilizing these structures. TMPyP4 can bind to all G4 topologies, however, compared to PDS, it exhibits lower specificity in discriminating G4 structures from double-stranded DNA (Ruan TL et al., 2017). Treatment of HeLa cells with increasing concentration of G4 ligands resulted in reduced cell survival under both normal and irradiated growth conditions. Based on literature survey and calculated IC_50_, the concentration of 5 µM PDS and 10 µM TMPyP4 were used to treat the cells for further experiments. Since both G4 ligands and gamma radiation are known to cause double strand breaks (DSB), the frequency of DSB formation was assessed at 30min and 2h post treatment. Immunofluorescence assay (IFA) revealed minimal or undetected γH_2_AX foci in untreated and in cells treated with PDS (5 µM) or TMPyP4 (10 µM) alone. However, a significant increase in γH_2_AX foci was observed at 30 min post irradiation in both untreated and G4 ligand treated cells (Fig. 1A, 1B). Analysis at 2 h post-irradiation showed that γH2AX foci levels returned to near-basal levels in untreated irradiated as well as in PDS-treated irradiated cells, indicating efficient repair of radiation-induced DNA damage (Fig. 1A, 1C). Surprisingly, the TMPyP4 treated irradiated cells did not exhibit comparable DNA damage repair kinetics. Although a reduction in γH_2_AX foci was observed at 2 hrs post irradiation, the foci count remained significantly above basal levels (Fig. 1A, 1C). Another notable observation from the IFA analysis was the lack of co-localization between G4 foci and DSB foci within the cells. This corroborates with the earlier reported observations (Kumari N et al., 2019).

**Figure 1.**
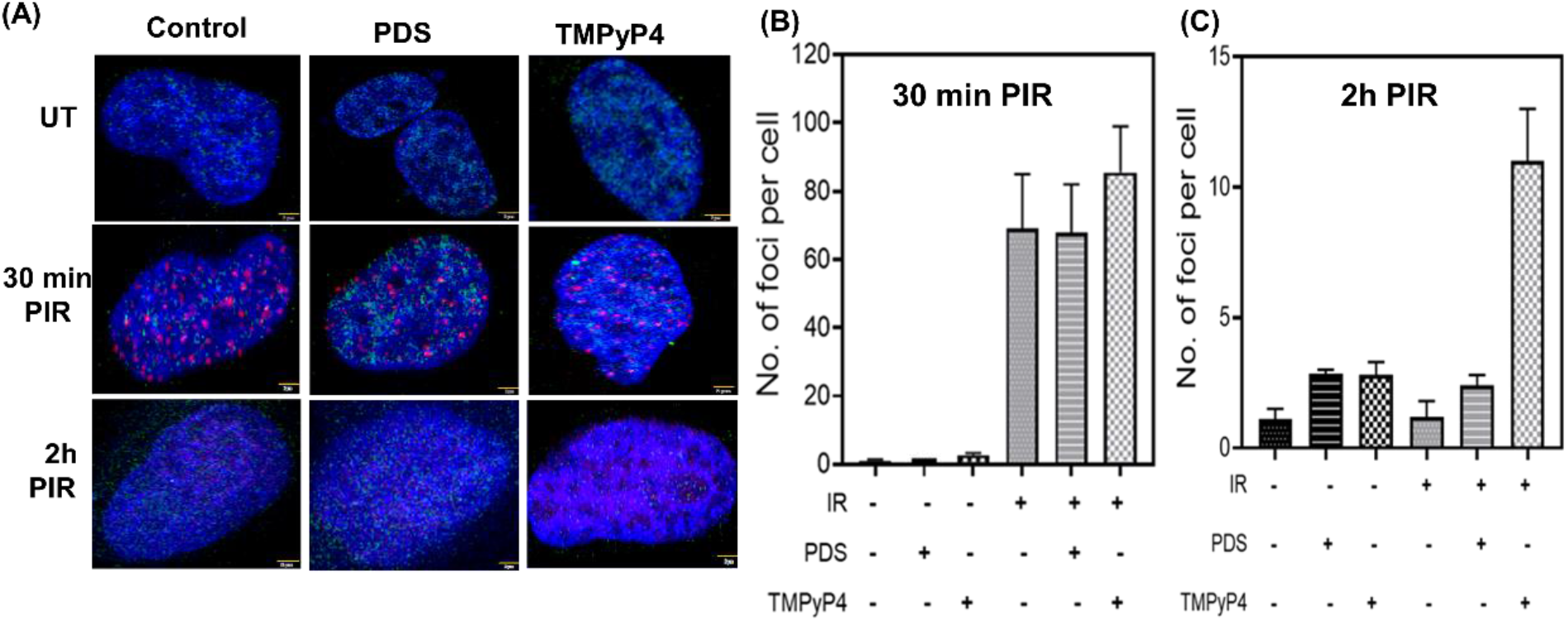
Effect of G4 binding ligands and ionizing radiation on DSB repair kinetics. **(A)** Immunofluorescence analysis of HeLa cells to assess double strand break repair kinetics. Cells were subjected to γ-radiation (8 Gy) either in the absence of G4 ligand (Control) or in the presence of PDS (5 µM) and TMPyP4 (10 µM). Post Irradiation Recovery (PIR) cells collected at 30 min and 120 min (2 h) were visualized for *in vivo* G4 structure formation using G4 structure-specific antibody (clone 1H6) and DNA repair kinetics using γH_2_AX antibody. Representative microscopic images are shown as merged channels (DAPI for nucleoid, Alexa fluor 488 for G4 structure and tetramethylrhodamine isothiocyanate (TRITC) for γH_2_AX). Scale bar 5 µm. Number of γH_2_AX foci per cell at **(B)** 30 min, and **(C)** 2h post irradiation was counted using automated CellSens software, and graphs were plotted using GraphPad Prism 8.

Next, the role of G4s in regulating small RNA expression under gamma radiation treated and untreated conditions was investigated using small RNA transcriptome analysis. The overall experimental workflow is illustrated in Fig. 2A. Total RNA was isolated from all six group of cells (H, HI, HP, HPI, HT, HTI) and sequenced as described in the Methods section (Fig. 2C). NextSeq Illumina reads generated an average of 30.4 million 75-bp single-end sequence reads per sample, ranging from 25.4 to 42.3 million sequence reads. More than 50 % of the Illumina reads exhibited PHRED scores above 30 (Q > 30) and were subsequently proceeded for downstream data analysis, resulting in an average of 11.2 million trimmed reads per sample. A variety of small RNA components were detected across the samples, such as miRNAs, rRNAs, tRNAs, snRNAs, snoRNAs, and piRNAs. The distribution of different small RNA species among the sample groups are depicted in Fig. 2B. On average, 984 known miRNAs were detected across all samples. Treatment with PDS and TMPyP4 differentially affected the abundance of known miRNAs. PDS treatment increased the proportion of detected known miRNA percent from 4% (in H, HI) to 4.5% (in HP, HPI). In contrast, TMPyP4 treatment decreased the proportion of detected known miRNA percent from 4% (in H, HI) to 3.29% (in HT) and 3.75% (in HTI). Further analysis revealed the expression of both novel and known miRNAs in all experimental groups. Although, a substantial number of known miRNAs exhibited read counts greater than 50, only a limited number of novel miRNAs showed read counts above this threshold. Therefore, further investigation was focussed primarily on known miRNAs (Fig. 2D, 2E).

**Figure 2.**
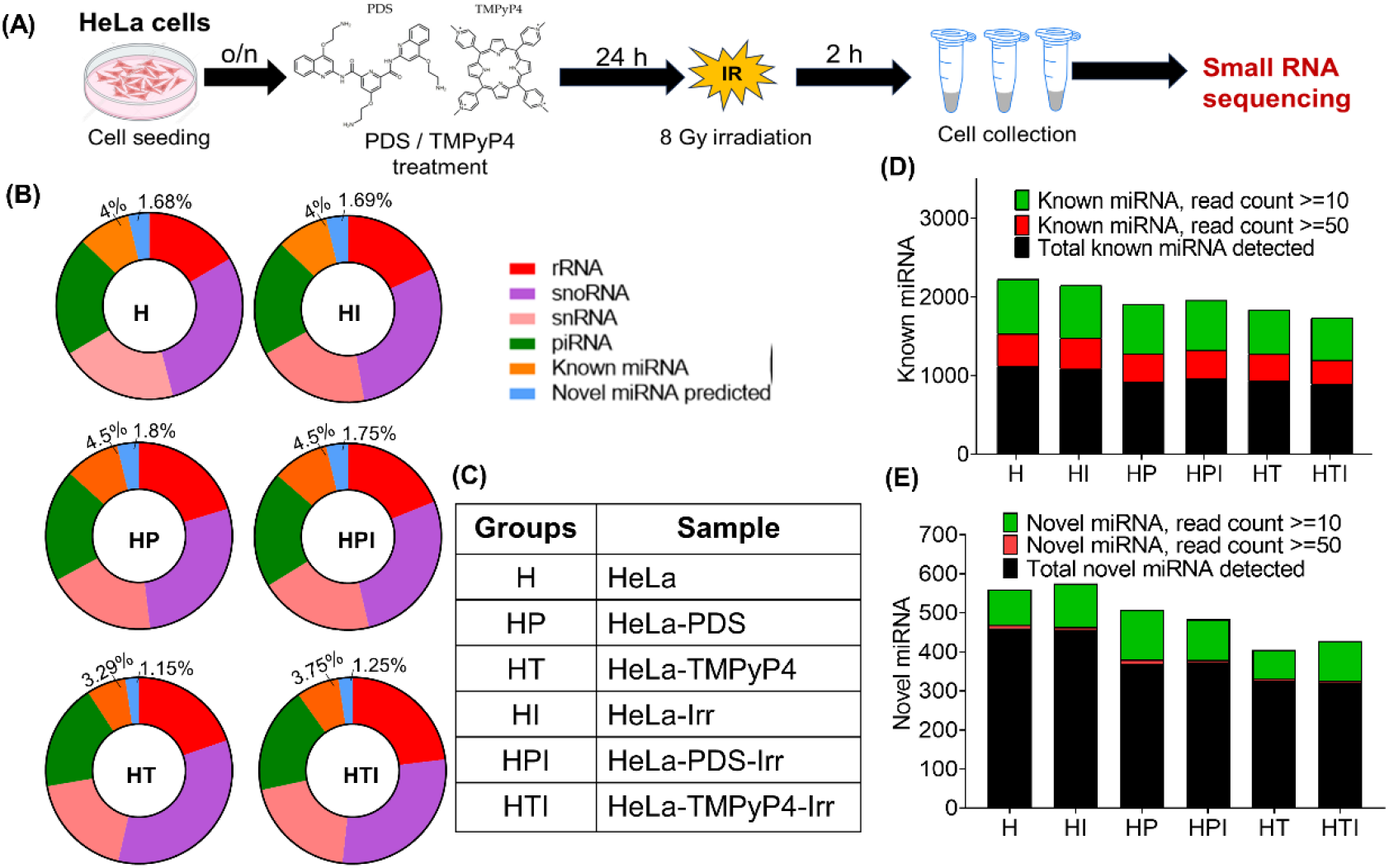
Small RNA transcriptome analysis to investigate the role of G4 structures (G4s) in small RNA expression under gamma radiation and G4 ligand treated conditions. **(A)** A schematic representation of the experimental set-up employed for small RNA sequencing. **(B)** Distribution of small RNA species in HeLa cells. The pie chart represents the proportion of small RNA molecules mapped to different types of RNAs. **(C)** Table representing different experimental groups and their respective samples. Differentially expression analysis of **(D)** known miRNAs and **(E)** novel miRNAs in cells treated with γ-radiation and G4 binding ligands PDS and TMPyP4.

### 3.2 Identification of differentially expressed known miRNAs across untreated and treated HeLa cells

Several specific miRNAs are known to regulate key genes involved in DNA damage response (Visser H and Thomas AD., 2020; Li Y et al., 2022). Since both G4 ligands treatment and gamma radiation results in DNA damage, changes in the expression profile of known miRNAs involved in DNA damage response as well as other cellular pathways were investigated. Differential expression analysis was performed by comparing untreated control cells with the different experimental groups after normalization of the sequencing data. The total number of differentially expressed miRNAs were 1012 (H vs HP), 1003 (H vs HT), 1170 (H v/s HI), 1105 (H v/s HPI), and 1054 (H v/s HTI). Among these, the number of statistically significant (p-value <0.05) with log_2_ fold change >1.5 were 16 (H vs HP), 23 (H vs HT), 32 (H v/s HI), 17 (H v/s HPI), and 125 (H v/s HTI). Volcano plots for each comparison are represented in Fig. 3A-3E. Several known miRNAs that are regulators of cell proliferation and cell cycle progression, such as miR-16-2-3p, miR-98-3p, miR-148b-3p, miR-16-2-3p, miR-125b-1-3p and miR-654-3p were differentially expressed across the treatment groups (Sati ISEE and Parhar I., 2021; Rodrigues DVS et al., 2018). In addition, multiple miRNAs targeting genes involved in DNA damage response pathways, such as miR-502, miR-101, miR-136, miR-34, miR-96, miR-26 and miR-27, also showed altered expression pattern (Visser H and Thomas AD., 2020; Li Y et al., 2022). Furthermore, miR-107, that is known to directly interact with Dicer, was found to be downregulated in HTI cells compared to control cells (Ristori E et al., 2015). Overall, the analysis revealed that the HTI group showed the highest number of differentially expressed miRNAs relative to control cells, with a substantial number of miRNAs being associated with the regulation of cell proliferation.

**Figure 3.**
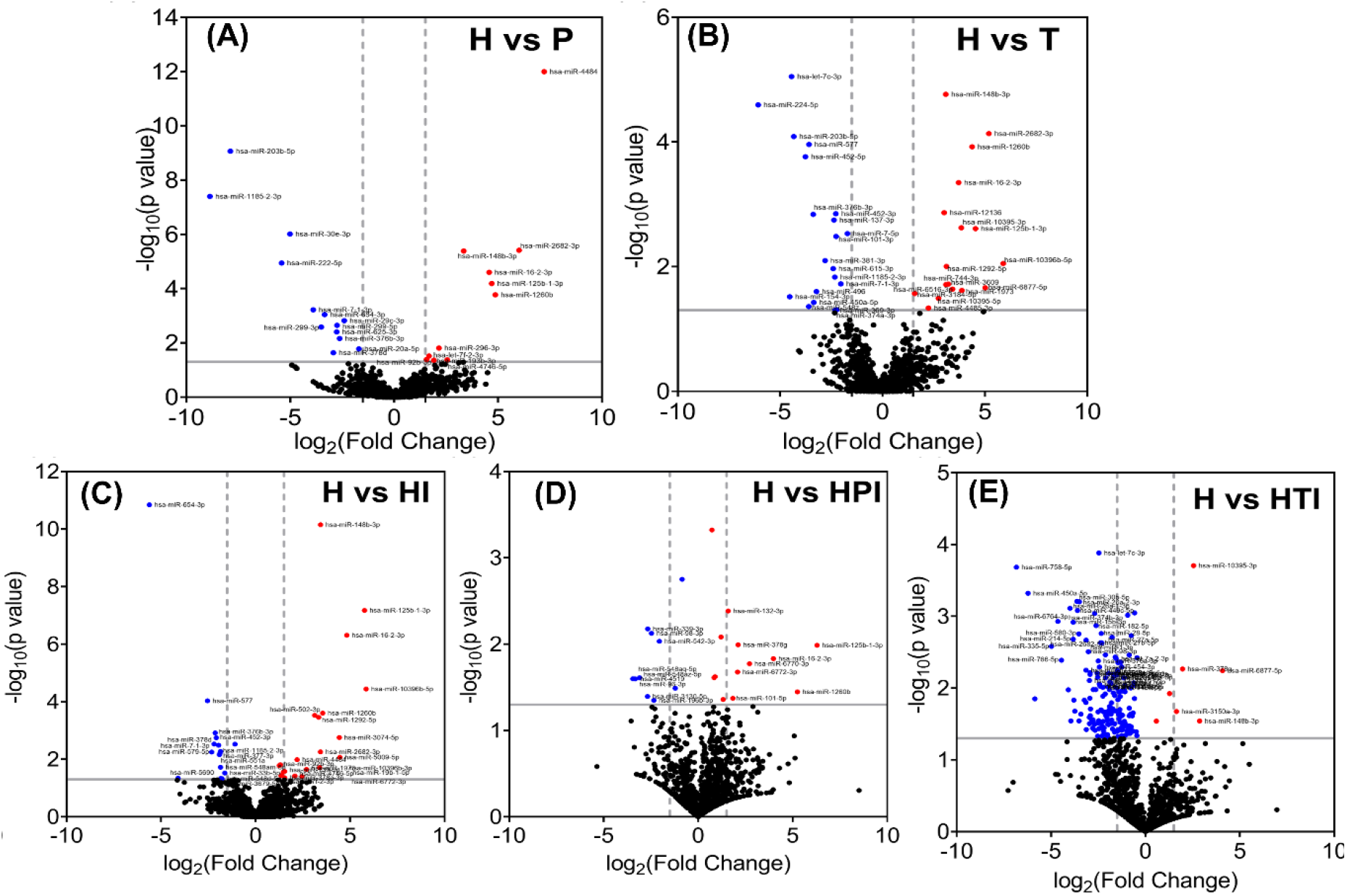
Volcano plots showing significantly upregulated and downregulated miRNAs under different treatment conditions: **(A)** HeLa (H) vs PDS treated (HP), **(B)** HeLa (H) vs TMPyP4 treated (HT), **(C)** HeLa (H) vs HeLa irradiated (HI), **(D)** HeLa (H) vs PDS treated and irradiated (HPI) and **(E)** HeLa (H) vs TMPyP4 treated and irradiated (HTI). The *P*-value cutoff for statistical significance is indicated by horizontal grey line while the vertical dashed grey lines at −1.5 and +1.5 indicate the log_2_ cutoff for significant fold change.

### 3.3 Analysis of gene ontology and KEGG pathways

Following the identification of differentially expressed miRNAs in each experimental group, target genes of the top 5 differentially expressing miRNAs (up and down) were predicted by TargetScanHuman 8.0. On the whole, HTI group exhibited the highest number of predicted target genes, consistent with the greater number of differentially expressed miRNAs identified in this group. To understand the biological processes modulated by these miRNAs, Gene Ontology (GO) and KEGG pathway enrichment analyses were also performed using the top 5 significantly upregulated and downregulated miRNAs from each group. Data analysis showed that majorly the pathways associated with signal transduction were affected as a result of differential miRNA expression arising from changes in cellular G4 dynamics. Several pathways like TGF-β (Transforming Growth Factor-beta) signalling, Hippo signalling, MAPK signalling, Wnt signalling, and Rap1 signalling were commonly enriched in all groups whereas RAS signalling pathway enrichment was specifically seen in the HI, HPI, and HTI groups. Notably, the MTOR signalling pathway which is primarily involved in cell growth, differentiation and metabolism, was enriched exclusively in the HTI group (Fig. 4A-4E). Similarly, GO analysis indicated that molecular functions pertaining to transcription factors DNA binding activity including RNA polymerase II transcription regulatory functions were affected across all the groups. Consistently, the biological pathways related to signal transduction or transcription regulation by RNA polymerase II or RNA processing were influenced in all the comparison groups. Thus, the overall analyses suggest that changes in G4 structure dynamics caused by Pyridostatin and TMPyP4 majorly affected the functions involved in chromatin remodelling and gene transcriptional regulation.

**Figure 4.**
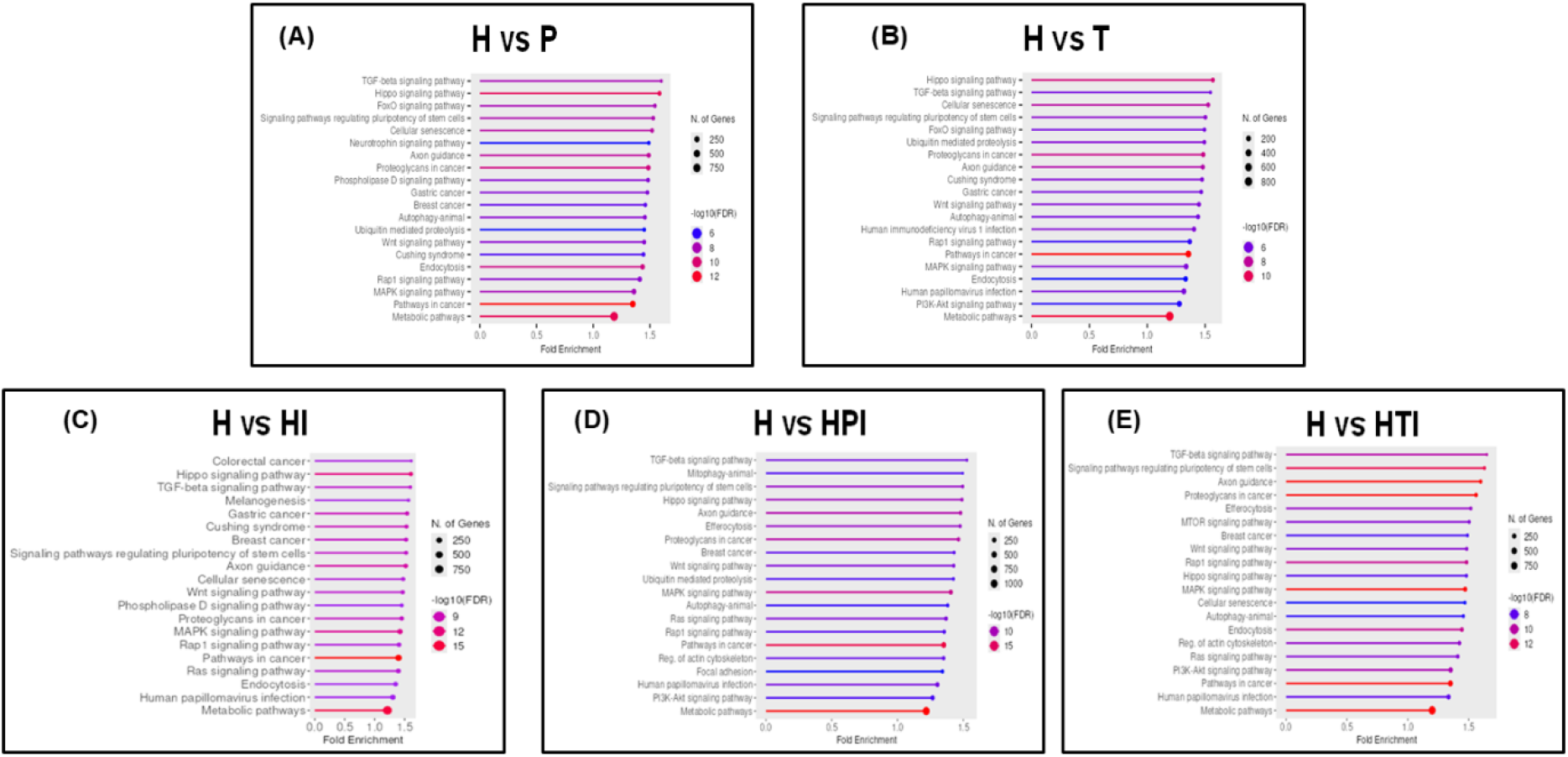
KEGG pathway enrichment analysis of the top 5 significantly upregulated and downregulated miRNAs from different treatment groups. **(A-E)** Using ShinyGO, the significantly differentially expressed miRNAs in each group were subjected to enrichment analysis. Enrichment bar plots for KEGG with FDR < 0.05.

### 3.4 The precursors of miRNA-4284 and miR-1226-3p fold into guanine quadruplex structures in vitro

G4 structure formation in pri-miRNAs and pre-miRNAs regulate their processing (Rouleau SG et al., 2018; Arachchilage MG et al., 2015). In the current study, a few top differentially expressed miRNAs harboured putative G4 forming motifs in their pre-miRNAs, for instance, the ones with FDR<0.05: let-7c-3p, miR-654-3p, miR-502-3p, miR-4484, miR-2682-3p, miR-1260b, miR-1292-5p, and miRNAs with p-value<0.05: miR-496, miR-12136, miR-378d, miR-551a. Among the earlier reported pre-miRNAs containing rG4 motifs, miR-92b was found to be upregulated in H and HP; miR-let-7 family was differentially regulated among multiple treatment cell groups; and miR-26a was found to be downregulated in the HTI group. Interestingly, miR-1260b and miR-551a harbors two putative G4 forming motifs. The miR-551a is differentially expressed in HI group while miR-1260b is upregulated in HI and HPI groups. Earlier, miR-1260b was shown to promote tumor growth in non-small cell lung cancer (NSCLC) and miR-551a has tumor suppressor role in breast cancer cells (Xia Y et al., 2019; Anuj et al., 2019). Furthermore, the formation of G4 structures in the pre-miRNAs of miR-4284 and miR-1226-3p; that are differentially expressed in HPI compared to HI was evaluated using circular dichroism (CD) and fluorescence-based assays. The results revealed that the pre-miRNAs of both miR-4284 and miR-1226-3p form G4 structures *in vitro*. This folding was demonstrated by an increase in the fluorescence intensities of ThT (Fig. 5A and 5B) in the presence of these pre-miRNA sequences compared to the dyes alone. Additionally, CD spectroscopy confirmed that these sequences fold into a parallel G4 topology (Fig. 5C and 5D), which is consistent with the bioinformatic analysis, where guanine-rich regions are predicted to form secondary G4 structures (Fig. 5E). Positive controls taken-G_2_LL and G_2_U also folded into parallel structures (Wieland and Hartig, 2007). However, the role of these intramolecular G4s within the pre-miRNAs of miR-4284 and miR-1226-3p on their function/processing needs to be investigated.

**Figure 5.**
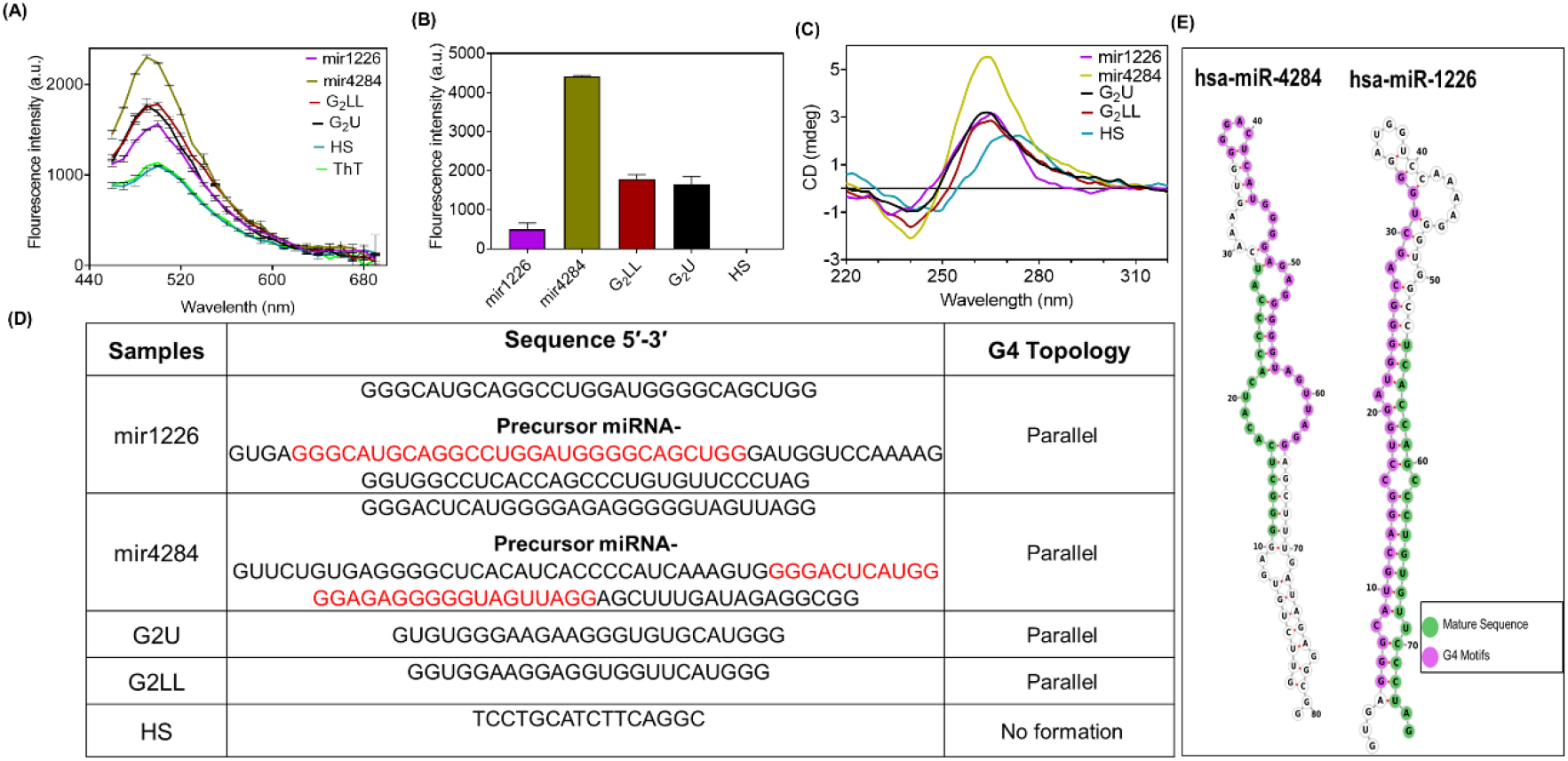
*In vitro* validation of G4 formation and bioinformatic secondary structure predictions of pre-miRNAs. The *in vitro* formation of G4 structures within the pre-miRNAs of hsa-miR-4284 and hsa-miR-1226-3p was evaluated using fluorescence-based assays and circular dichroism (CD) spectroscopy. Thioflavin T (ThT) assay: Annealed sequences were incubated with 0.5 µM ThT. **(A)** Fluorescence emission spectrum recorded from 450 nm to 700 nm with a fixed excitation at 425 nm. **(B)** Comparison of fluorescence intensities recorded at an emission of 490 nm following excitation at 425 nm. G_2_LL and G_2_U were taken as positive control while HS as negative control. All graphs were plotted using GraphPad Prism 8.0. **(C)** CD spectra of the putative G4-forming motifs within pre-hsa-miR-4284 and pre-hsa-miR-1226-3p in the presence of 50 mM KCl. **(D)** Summary table showing putative G4-forming motifs folding into distinct parallel G4 topologies. **(E)** Computational secondary structure models of the pre-miRNA stem-loop hairpins.

### 3.5 Putative G4 DNA motifs at 5’ and 3’ untranslated regions of histone deacetylases and methyl transferases fold into stable G4s

Stabilization of G4s with small-molecule ligands has previously been shown to change DNA and histone modifications (Mukherjee AK et al., 2019). In addition, previous reports have revealed that post translational modification of DGCR8, particularly through deacetylation, can modulate miRNA processing (Wada T et al., 2012). Histone deacetylases (HDACs) and methyl transferases are the major enzymes involved in histone modifications. The expression of some of the enzymes of these classes in response to PDS and TMPyP4 treatment was studied. The 5′ and 3′ UTRs histone deacetylases and methyl transferases were searched for the presence of putative G4 sequences. The ability to form stable G4 structures by some of the selected sequences of HDAC1, HDAC2, SETD7, and SUV39H1 which showed high G-scores, were evaluated in vitro using circular dichroism spectroscopy (Fig. 6B). As shown in the figure, the results demonstrated that the identified motifs fold into various topologies like parallel, hybrid and anti-parallel G4 conformation (Fig. 6A, 6B). Subsequently, the effect of PDS and TMPyP4 on the expression levels of HDAC1, HDAC2, SETD7, and SUV39H1 was investigated by RT-qPCR. The results revealed that PDS treatment reduced the transcript levels of HDAC1 and HDAC2 whereas the transcript levels of SETD7 and SUV39 levels were upregulated. Similar results were observed with TMPyP4 treatment except the SUV39H1 levels were down regulated (Fig. 6C). Thus, these findings suggest that G4 dynamics plays a significant role in modulating the expression of histone modifying enzymes in HeLa cells, at least for the enzymes tested in this study.

**Figure 6.**
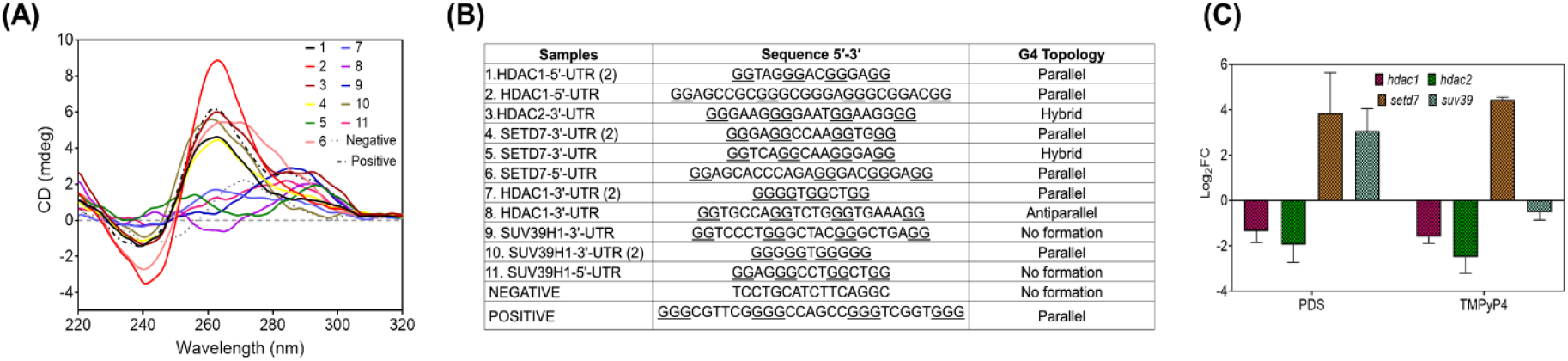
In vitro characterization of G-quadruplex (G4) structure formation by putative G4 forming sequences (PQS) present at untranscribed regions (UTRs) of histone deacetylases (HDAC1 & HDAC2) and methyl transferases (SETD7 & SUV39H1) along with real time PCR analysis. **(A)** Circular dichroism (CD) spectra of selected PQS from the UTRs of HDAC1, HDAC2, SETD7, and SUV39H1, in the presence of 50 mM KCl demonstrating their folding into distinct G4 topologies **(B)** Summary table showing PQS details and their topology obtained in the presence of KCl. **(C)** Relative transcript expression levels of HDAC1, HDAC2, SETD7, and SUV39H1 measured via RT-qPCR following treatment with G4-stabilizing ligands PDS and TMPyP4. Glyceraldehyde-3-phosphate dehydrogenase (GAPDH) was used as endogenous control for normalising mRNA expression levels.

## 4. Discussion

Guanine quadruplexes regulate a number of biological processes depending on their location. Several factors, such as monovalent cation concentration, presence of single strand DNA and G4 interacting proteins, influence the in vivo G4 structure dynamics (Zhang et al., 2023). Small planar aromatic molecules bind these structures with high affinity, hinder their natural folding-unfolding process, and perturb the G4 regulated cellular processes (Romano F et al., 2025). Two widely studied G4 stabilizing ligands are-Pyridostatin and TMPyP4. PDS, a heterocyclic compound, is highly selective for G4 structures and induces replication stress, DNA damage and severe transcriptional changes (Groelly FJ et al., 2022). While TMPyP4, is a porphyrin-based non selective G4 ligand that is mainly used to study telomere dysfunction (Grand CL et al., 2002). In this study, we investigated how these two G4 ligands influence the genome-wide expression of microRNAs under gamma radiation induced stress in HeLa cells. miRNAs are a class of small non coding RNAs that regulate post-transcriptional gene expression. Under the condition used for small RNA analysis, PDS (5 µM; 24 h) and TMPyP4 (10 µM; 24 h) in combination with 8 Gy gamma radiation, the HeLa cell viability remained unaffected (Fig. 1). Previously, it was reported that HeLa and MCF7 cells treated with PDS (2 μM; 5 h), or TMPyP4 (5 μM; 5 h), followed by 10 Gy radiation showed fewer gamma radiation induced double strand break (DSB) foci compared to irradiated control cells, suggesting intracellular G4 stabilization may protect the genome (Kumari N et al., 2019). However, in this study, TMPyP4 treated cells showed a higher number of gamma radiation induced DSB foci at 2 h after radiation compared to irradiated control cells. This difference in observations could be attributed to variations in the concentration and duration of ligand treatment between the studies.

In the present investigation, the total miRNA content (in percent) increased in PDS treated cells while decreased in cells exposed to TMPyP4 (Fig. 2). The highest number of differentially expressed miRNAs was observed in the HTI group (TMPyP4 + gamma radiation). As both PDS and TMPyP4 interact with DNA and RNA G4 structures, global changes in miRNA expression profiles can be anticipated due to the involvement of these biomolecules in miRNA biogenesis. The differing effects of the ligands may arise due to their distinct affinity and selectivity towards G4 structures, which might have caused the observed differences. TMPyP4 has been reported to destabilize the RNA quadruplexes (rG4s), changing the equilibrium between G4 and stem-loop structures in precursor miRNAs and thereby affecting the miRNA genesis. For instance, miRNA-149 and miR-1587 expression were elevated as a result of destabilization of pre-miRNA by TMPyP4 (Ghosh A et al., 2019; Li et al., 2023). In some cases, TMPyP4 was shown to aggregate rG4 structures and hinder their interaction with proteins such as that between let-7e miRNA with nucleolin (Santos T et al., 2022). In contrast, PDS is a potent stabilizer of rG4, and down regulates RNA binding proteins in HeLa cells (Hou Y et al., 2022). In addition, cellular stress promotes and stabilizes G4 structures protecting the nucleic acids from degradation (Kharel et al., 2023; Tewari et al., 2025). Thus, increased G4s stability in presence of PDS and gamma radiation may contribute to the rise in total miRNA content observed in HPI cells, a possibility which will be addressed separately.

RNA G4 forming sequences have previously been reported within some primary miRNAs (pri-miRNAs) (Rouleau SG et al., 2018). In the present investigation, out of the reported pri-miRNAs containing putative rG4 sequences, pri-miR-497 was found to be downregulated in the HTI group while pri-miR-451a was downregulated in the HPI group. Further, it has been demonstrated that around 16% of all identified human precursor-miRNAs (pre-miRNas) sequences harbour G4 motifs which can form stable G-quadruplex structures instead of canonical stem loops, directly affecting mature miRNA formation (Ghafouri-Fard S et al., 2022; Tassinari et al., 2021; Arachchilage MG et al., 2015). This current study revealed that pre-miRNAs of hsa-miR-4284 and hsa-miR-1226-3p are folding into parallel structures and are differentially regulated among different groups. These findings further support the notion that G4 stabilization using ligands can alter miRNA biogenesis through the presence of G4 structures within primary and precursor miRNA sequences. To the best of our knowledge, this study is the first to report regulation of intracellular miRNA levels under combined conditions of gamma radiation stress and G4 stabilization in HeLa cells.

Histone modifications are crucial parts of epigenetic regulation of gene expression. In this report, it was observed that 5’ and 3’ UTR of genes encoding for histone modifiers harbour putative G4 forming sequences which fold into G4 structures in vitro (Fig. 6A, 6B). A complex reciprocal regulatory network exists between histone modifications and miRNA processing - like histone modifications determine the chromatin accessibility for miRNA genesis and miRNA can post transcriptionally regulate the histone modifying enzymes expression (Szczepanek J and Tretyn A., 2023). Under these conditions, the reduced deacetylase levels in G4 ligand treated cells (Fig. 6C) may have caused hyper acetylation of histones resulting in open chromatin, leading to increased miRNA biogenesis, a possibility that could not be ruled out. Furthermore, the observed effect on miRNA profile may also result from combinatorial histone modifications.

Radiotherapy is widely used in cancer care (over >60% cases), where it is given either to treat symptoms or to destroy tumors. It is often used alongside surgery or chemotherapy or immunotherapy to maximize the treatment outputs (Elbanna M et al., 2021; Yu S et al., 2022). Recently, the therapeutic potential of G4 ligands in cancer cells is also explored as they can effectively inhibit telomere replication or oncogene expression (Iachettini S et al., 2024). To explore the possibility of combining these two treatment modalities, as well as to develop more effective and personalized medicine, a thorough understanding of the affected molecular pathways is required. Towards this goal, the present study has first time reported how the G4 ligands-PDS and TMPyP4, change the global miRNA profile under radiation stress. Exploring the possibility of targeting these differentially expressed miRNAs under gamma radiation treatment conditions may further increase the treatment response in cancer cells.

## Acknowledgment

We sincerely acknowledge Dr. Hema Rajaram, Bhabha Atomic Research Centre, Mumbai, for her constant support and suggestions during this work. Ms Himani Tewari and Ms Sindhuja Singh express their gratitude to the Department of Atomic Energy, Government of India, for their fellowship. We sincerely thank the Department of Atomic Energy for financial support.

## Authors’ contributions

SM conducted the experiments, analysed the data, and wrote the manuscript. HT analysed the data and wrote the manuscript. SS performed the experiments. SK conceptualized the idea, analysed the data, wrote the manuscript, and communicated.

## Funding

This research is supported by the Department of Atomic Energy, Government of India.

## Conflict of interest statement

The authors declare no conflict of interest.

## Notes

### Competing Interest Statement

The authors have declared no competing interest.

